# Delivery of novel replicating vectors to *Synechococcus sp.* PCC 7002 via natural transformation of plasmid multimers

**DOI:** 10.1101/2024.07.31.606084

**Authors:** Cody Kamoku, Cheyanna Cooper, Ashley Straub, Nathan Miller, David R. Nielsen

## Abstract

In most cyanobacteria, genetic engineering efforts currently rely upon chromosomal integration; a time-consuming process due to their polyploid nature. To enhance strain construction, here we develop and characterize two novel replicating plasmids for use in *Synechococcus* sp. PCC 7002. Following an initial screen of plasmids comprising seven different origins of replication, two were found capable of replication: one based on the WVO1 broad host range plasmid and the other a shuttle vector derived from pCB2.4 from *Synechocystis* sp. PCC 6803. These were then used to construct a set of new replicating plasmids, which were shown to be both co-transformable and stably maintained in PCC 7002 at copy numbers between 0.6-1.4 and 7-16, respectively. Lastly, we demonstrate the importance of using multimeric plasmids during natural transformation of PCC 7002, with higher order multimers providing a 30-fold increase in transformation efficiency relative to monomeric plasmids. Useful considerations and methods for enhancing multimer content in plasmid samples are also presented.

## 1. Introduction

An inherent challenge to cyanobacterial engineering stems from the polyploid nature of their chromosomes. *Synechococcus* sp. PCC 7002 (henceforth PCC 7002), for example, maintains 5-12 copies of its chromosome [1]. This can be problematic since the ability to fully knock out a gene or stably insert one requires complete chromosomal segregation. This typically requires a lengthy process of repeated restreaking on plates with increasing antibiotic concentrations; an especially arduous task if the resulting edit is deleterious to strain fitness. Although this requirement could be bypassed via the use of replicating episomal plasmids, to date, the only broad host range plasmids shown to replicate in PCC 7002 include RSF1010 [2] and the newly discovered pYS from *Synechocystis* sp. PCC 6803 (henceforth PCC 6803) [3]. Meanwhile, although several native cyanobacterial plasmids have also been repurposed as shuttle vectors (including pANS from *Synechococcus elongatus* PCC 7942 [4] and pCA2.4, pCB2.4, pCC5.2 from PCC 6803 [5–7]), their use has so far remained limited to the strain from which they were derived and not yet extended to PCC 7002.

Wide-spread adoption of replicating plasmids in cyanobacteria is further hindered by a lack of robust transformation methods. Traditional delivery methods predominantly rely upon conjugation [2] or, less commonly, electroporation [8]. While conjugation is efficient, the process is time consuming and poorly suited for co-transforming multiple constructs [9]. Electroporation, on the other hand, requires large amounts of DNA as it suffers from low efficiencies; possibly due to the exopolysaccharide layer that covers cyanobacterial cells [10]. Another method of replicating plasmid delivery exploits the native capability of naturally transformable bacteria to uptake multimeric plasmids which are then reassembled into their monomeric form once inside the cell. Specifically, plasmid multimers (which can be generated via in vitro methods [11] and often occur naturally within samples miniprepped from *Escherichia coli* [12,13]) are imported as ssDNA and may become recircularized if: a) they contain a self-priming mechanism to direct the cell to synthesize the complementary strand of DNA, such as a ssDNA origin of replication [14] or a primosome assembly site [15]; or b) complementary ssDNA is imported and serves as a primer for dsDNA synthesis [15]. While replicating plasmid delivery via multimeric plasmids has been demonstrated for a diverse range of naturally transformable bacteria (including *Bacillus subtilis* [11], *Acinetobacter baylyi* ADP1 [16], and several *Vibrio* species [17]) and archaea (including *Thermococcus kodakarensis* KOD1 and *Pyrococcus yayanosii* A1 [18]), to our knowledge, such methods have yet to be explicitly studied or applied in cyanobacteria.

## 2. Materials and methods

### 2.1 Strains, media and culture conditions

All strains constructed and/or used in this study are summarized in Table 1. *E. coli* DH5α, DH10β, and DHIP1 were grown in LB broth at 37^°^C. *E. coli* DHIP1 is derived from DH10β and expresses *slr0214* from PCC 6803 (encoding a DNA methyltransferase targeting the HIP1 sequence: GCGATCGC) [19] from the *gidB-atpI* intergenic locus. PCC 7002 strains were routinely cultured using media A+ [20] with 150 μE light intensity under a 1% CO_2_ atmosphere. kanamycin (75 μg/mL) and/or gentamicin (30 μg/mL) were used to select for and maintain replicating plasmids in PCC 7002.

**Table 1.** Strains and plasmids constructed and/or used in this study.

| Strain | Relevant Characteristics |
| --- | --- |
| *E. coli* DH5α | *mrr-hsdRMS-mcrBC* |
| *E. coli* DH10β | $\Delta(mrr-hsdRMS-mcrBC)$ |
| *E. coli* DHIP1 | DH10β *atpi:slr0214:gidB* |
| *Synechococcus* sp. PCC 7002 | Wild type |
| Plasmid | Relevant Characteristics |
| pCMK74 | WVO1 ori, *Kan*<sup>R</sup> |
| pCMK191 | WVO1 ori, *Gm*<sup>R</sup> |
| pCMK139 | pSC101 ori (RepA<sup>N99D</sup> $\Delta par$),<br>pCB2.4 ori, *Kan*<sup>R</sup> |
| pCMK188 | pBR322 ori, pCB2.4 ori, *Kan*<sup>R</sup> |

### 2.1 Plasmid and strain construction

Plasmid pBAV1K-T5-gfp (a gift from Prof. Ichiro Matsumura; Addgene #26702) [21], was used to construct all plasmids containing the WVO1 origin of replication (i.e., pCMK74 and pCMK191). To construct pCMK74, the GFP sequence was first removed. Next, between the kanamycin resistance gene and the WVO1 origin of replication, a HIP1 site was added since its methylation has been reported to improve transformation efficiency in other cyanobacteria [19]. To construct pCMK191, the kanamycin resistance gene in pCMK74 was replaced with a gentamicin resistance cassette. Plasmid pSL316 (a gift from Prof. Himadri Pakrasi) [5], was used to construct all pCB2.4-derived shuttle vectors (i.e., pCMK139 and pCMK188). To construct pCMK139, the streptomycin resistance gene and GFP cassette in pSL3136 were first replaced with a kanamycin resistance gene. To increase plasmid copy number, the par locus of pSC101 was then deleted and the RepA gene mutated (RepA^N99D^). To construct pCMK188, the pSC101 origin in pCMK139 was replaced with the pBR322 origin of replication. A HIP1 site was likewise included between the kanamycin resistance gene and pCB2.4 origin of replication in both pCMK139 and pCMK188. Plasmid pBb(RSF1010)1k-GFPuv (a gift from Prof. Brian Pfleger; Addgene #106395) [22], was used as a positive control for natural transformation of all replicating plasmids. To improve comparison across plasmids, a kanamycin resistance cassette was first incorporated into each of pAM4788, pSL3136, and pSL3137. All DNA fragments were amplified by PCR using Q5 High Fidelity Polymerase (New England Biolabs; NEB). Plasmids were constructed via Golden Gate assembly [23] or Gibson Assembly [24]. As necessary, plasmids were sequenced via Oxford nanopore sequencing (Plasmidsaurus) to confirm sequence identity and determine their multimer content. Plasmids pCMK74, pCMK191, pCMK139, and pCMK188 have been submitted to and made available via Addgene.

### 2.3 Generation of plasmid multimers

Plasmid multimers were naturally generated by and obtained in miniprepped samples from *E. coli* DH5α, DH10β, or DHIP1. Alternatively, plasmid multimers were also synthesized from monomeric templates using the TempliPhi Phi29 Polymerase Amplification Kit (Cytiva). In this case, samples were purified via ethanol precipitation, after which the entire purified reaction was used to transform PCC 7002.

### 2.4 Generation of plasmid samples enriched with differently sized multimers

Plasmid samples enriched with differently sized multimers were resolved from miniprepped samples via gel electrophoresis (0.5% agarose) followed by extraction of individual bands using the QIAquick Gel Extraction Kit (Qiagen) and DNA Clean & Concentrator Kit (Zymo Research). Each isolated fraction was then analyzed via Nanopore sequencing (Plasmidsaurus) to determine the size and distribution of included plasmid multimers, if any.

### 2.5 Natural transformation of PCC 7002

Natural transformation of replicating plasmids into PCC 7002 was performed as follows. PCC 7002 was grown overnight at 37^°^C in 15 mL media in 250 mL shake flasks with 150 μE light intensity under a 1% CO_2_ atmosphere. The next morning, by which time culture had grown to an OD_730_ of 0.5-0.7, cells were collected by centrifugation at 3000 x *g*, after which the pellet was resuspended in fresh media A+ to an OD_730_ of 1.0. Next, 100-1000 ng of plasmid DNA was added to 1 mL of resuspended cells in a 15 mL culture tube, then incubated for 3 hours with shaking. Cells were then plated and incubated at 37^°^C with 150 μE light intensity and atmospheric CO_2_ for 6-7 days before the resulting colonies were counted.

### 2.6 Plasmid copy number determination by qPCR

Three independent colonies of PCC 7002 containing each plasmid were grown overnight to an OD_730_ of 0.5-0.7, at which point cells were harvested and total DNA was extracted using a Quick-DNA Fungal/Bacterial Miniprep kit (Zymo Research). qPCR was performed using a Quantstudio 3 (Thermo Fisher) and Forget-Me-Not EvaGreen qPCR Master Mix with low ROX (Biotium) according to manufacturer instructions. All qPCR primers were custom designed and synthesized by IDT (Table 2). Primers 7002_ppc_F and 7002_ppc_R target the chromosomal *ppc* locus [25], which was used as a reference for calculating plasmid copy number relative to chromosome copy number. Primer pairs ck925 and ck926 amplify the *repA* in WVO1-derived plasmids while ck927 and ck928 amplify *ORF3* in pCB2.4-derived shuttle vectors. Relative plasmid copy number was determined by dividing the moles of the plasmid amplicon by moles of the chromosomal amplicon, where moles were determined using a standard curve prepared for each PCR product [26]. Standards were generated by separately amplifying then gel purifying the same amplicons using either wild-type PCC 7002 genomic DNA or miniprepped plasmids as templates and used to generate standard curves. All biological samples and standard curve samples were run in technical triplicate. AccuBlue dsDNA Quantitation Kit (Biotium) was used to quantify DNA concentration in all samples and standards.

**Table 2.**
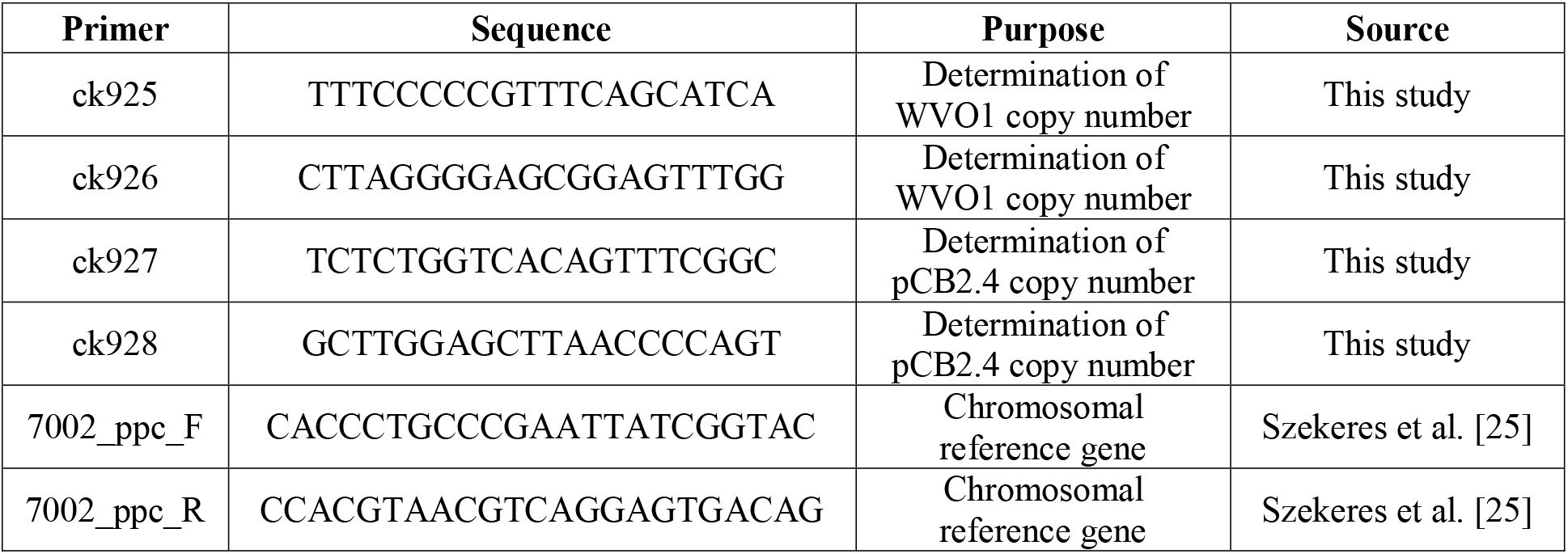
qPCR primers designed and used for copy number determination.

### 2.7 Plasmid stability and curing

Plasmid-harboring PCC 7002 strains were grown overnight in a 125 mL flask containing 25 mL media A+ with 75 μg/mL kanamycin. The next day, cells were inoculated into 25 mL fresh media A+ without antibiotics at an initial OD_730_ of 0.05. Cells were grown for 48 hours at 37^°^C while shaking at 220 rpm with 150 μE light intensity and 1% CO_2_, at which point a sample from each culture was serially diluted (10^−2^ to 10^−5^) then plated on 0.75% agar plates either with (75 μg/mL) or without kanamycin. After two days growth at 37^°^C with 150 μE light intensity and atmospheric CO_2_, total colonies were counted on both plates. At the same time, each culture was also used to inoculate a new flask containing 25 mL fresh media A+ without antibiotics at an initial OD_730_ of 0.05. This entire process was repeated for a total of three serial transfers.

## 3. Results

### 3.1 Screening diverse plasmid origins for replication in PCC 7002

We began by screening a pool of plasmids with diverse origins of replication to determine their propensity for replication in PCC 7002 (Table 3). Except for the two broad host range origins (i.e., WVO1 and pBBR1), most other candidates were chosen since they had previously been shown to replicate in at least one other cyanobacterial species (but not PCC 7002). RSF1010 served as a positive control since it is known to replicate and be stably maintained in PCC 7002 [27,28]. To ensure that all samples would contain multimers, the TempliPhi Amplification Kit was first used to generate multimeric copies of each plasmid via rolling circle amplification, after which the resulting mixtures were transformed into PCC 7002. Among all tested plasmids, only those with RSF1010, WVO1, or pCB2.4 origins yielded healthy colonies, whereas no colonies were obtained for the pBBR1 broad host range origin. Meanwhile, for each of the RK2, pANS, pDU1, and pCA2.4 origins, only very small colonies appeared several days after transformation, all of which were ultimately found to be non-viable upon being transferred to liquid culture.

**Table 3.** *Plasmids screened for natural transformation and replication in PCC 7002*. All plasmids contain *Kan*^*R*^. * indicates existing antibiotic resistance cassette was replaced with *Kan*^*R*^.

| Plasmid | Origin of replication | Attributes or Source | Replication in cyanobacteria | Current Result in PCC 7002 | Reference |
| --- | --- | --- | --- | --- | --- |
| pBb(RSF1010)1k-GFPuv | RSF1010 | Broad host range | PCC 6803, PCC 7002, PCC 7942 | Healthy colonies | Cook et al. [22] |
| pBAV1K-T5-gfp | WVO1 | Broad host range | Unknown | Healthy colonies | Bryksin et al. [21] |
| pSEVA221 | RK2 | Broad host range | PCC 6803 | Small colonies that quickly died | Martinez-Garcia et al. [29,30] |
| pBBR1MCS-2 | pBBR1 | Broad host range | Unknown | No colonies | Kovach et al. [31] |
| pAM4788* | pANS | *Synechococcus elongatus* sp. PCC 7942 | PCC 7942, *Anabaena* sp. PCC 7120. | Small colonies that quickly died | Chen et al. [4] |
| pRL2960 | pDU1 | *Nostoc* sp. PCC 7524 | PCC 7524 | Small colonies that quickly died | Wolk et al. [32] |
| pSL3136* | pCB2.4 | *Synechocystis* sp. PCC 6803 | PCC 6803 | Healthy colonies | Liu and Pakrasi [5] |
| pSL3137* | pCA2.4 | *Synechocystis* sp. PCC 6803 | PCC 6803 | Small colonies that quickly died | Liu and Pakrasi [5] |

### 3.2 Construction and direct transformation of novel replicating plasmids in PCC 7002

Since both pBAV1K-T5-gfp and pSL3136 showed promising initial results with respect to replication in PCC 7002, their associated origins of replication (WVO1 and pCB2.4, respectively) were next used to develop a set of novel replicating vectors for use in PCC 7002 WVO1 is a broad host range origin of replication [21] that, to our knowledge, has not previously been investigated in cyanobacteria. Meanwhile, pCB2.4 is derived from PCC 6803 and has previously been used to create a shuttle vector for that strain [5,7], but not yet for PCC 7002. Using these origins, two novel replicating plasmids were initially constructed: pCMK74 and pCMK188 (Figure 1).

**Figure 1.**
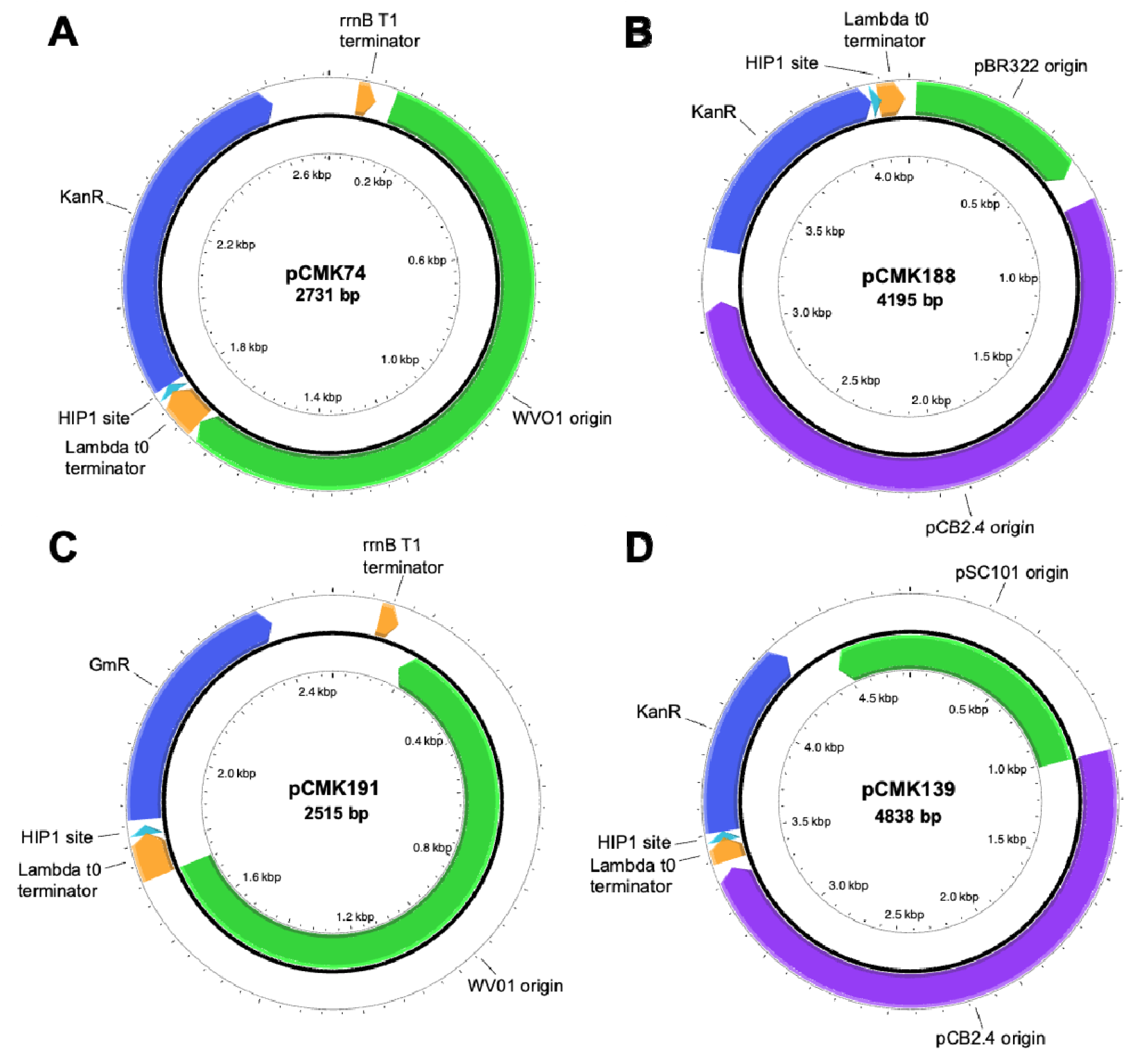
Plasmid maps representing the four novel plasmids constructed in this study. A: pCMK74, containing the broad host range WVO1 origin and Kan^R^. B: pCMK188, a shuttle-vector containing both pCB2.4 and pBR322 origins and Kan ^R^. C: pCMK191, containing the broad host range WVO1 origin and Gm ^R^. D: pCMK188, a shuttle-vector containing both the pCB2.4 origin and a mutated pSC101 origin and Kan ^R^. Plasmid maps made using Plasmapper 3.0 [34].

Following construction, pCMK74 and pCMK188 were miniprepped from *E. coli* DHIP1 and transformed into PCC 7002, after which the copy number of each during exponential growth was determined by qPCR. The number of plasmids per chromosome were measured as 1.48 ± 0.71 for pCMK74 and 0.13 ± 0.04 for pCMK188. For reference, in PCC 6803, the copy number of pCB2.4 per chromosome has been measured as 0.4 during exponential growth [33]. Assuming an average chromosome copy number of 5-11 for PCC 7002 [1], this suggests the plasmid copy number per cell is between 7-16 and 0.6-1.4 for pCMK74 and pCMK188, respectively. For comparison, in PCC 6803, more commonly employed RSF1010-derived plasmids have been reported to be maintained at around 31 ± 5 copies per cell, while RK2 plasmids are maintained at 9 ± 2 copies per cell [30].

### 3.3 Characterizing plasmid integrity, stability, and curability in PCC 7002

To confirm that they are indeed maintained as stable, episomal plasmids, pCMK74 and pCMK188 were next each miniprepped from PCC 7002, retransformed back into *E. coli*, and then miniprepped once more. Those samples were then subject to nanopore sequencing, from which complete sequence integrity of both plasmids was indeed confirmed. Next, to further explore the versatility of WVO1- and pCB2.4-derived plasmids as genetic engineering tools, their ability to be co-transformed and compatibly maintained together in PCC 7002 was investigated. To do so, however, the kanamycin resistance gene in pCMK74 was first replaced with a gentamicin resistance cassette, resulting in pCMK191 (Figure 1). pCMK191 and pCMK188 were then sequentially transformed into PCC 7002 and ultimately plated on media A+ agar plates containing both kanamycin and gentamicin. Co-transformed colonies were likewise enriched in liquid medium, after which both plasmids were miniprepped together from PCC 7002 and retransformed back into *E. coli. E. coli* transformants were selected on LB agar plates containing either kanamycin or gentamicin to enable individual isolation of each plasmid. Once again, nanopore sequencing revealed that the sequence integrity of both plasmids was fully maintained.

We lastly sought to characterize the curability of pCMK74 and pCMK188 in PCC 7002, as measured by their ability to persist in the absence of antibiotic selection. Cultures of PCC 7002 harboring each plasmid were grown without antibiotics over a total period of eight days. During that time, every second day, a sample from each culture was plated onto media A+ agar with and without antibiotics and serially transferred into fresh media. As seen in Figure 2, pCMK188 was much more stable than pCMK74; with approximately 24% of cells still retaining pCMK188 by day 8 versus just 0.09% for pCMK74. That said, for applications in which plasmid curability is desired (e.g., removal of Cas9 after CRISPR/Cas9 genome engineering), pCMK74 may prove most useful.

**Figure 2.**
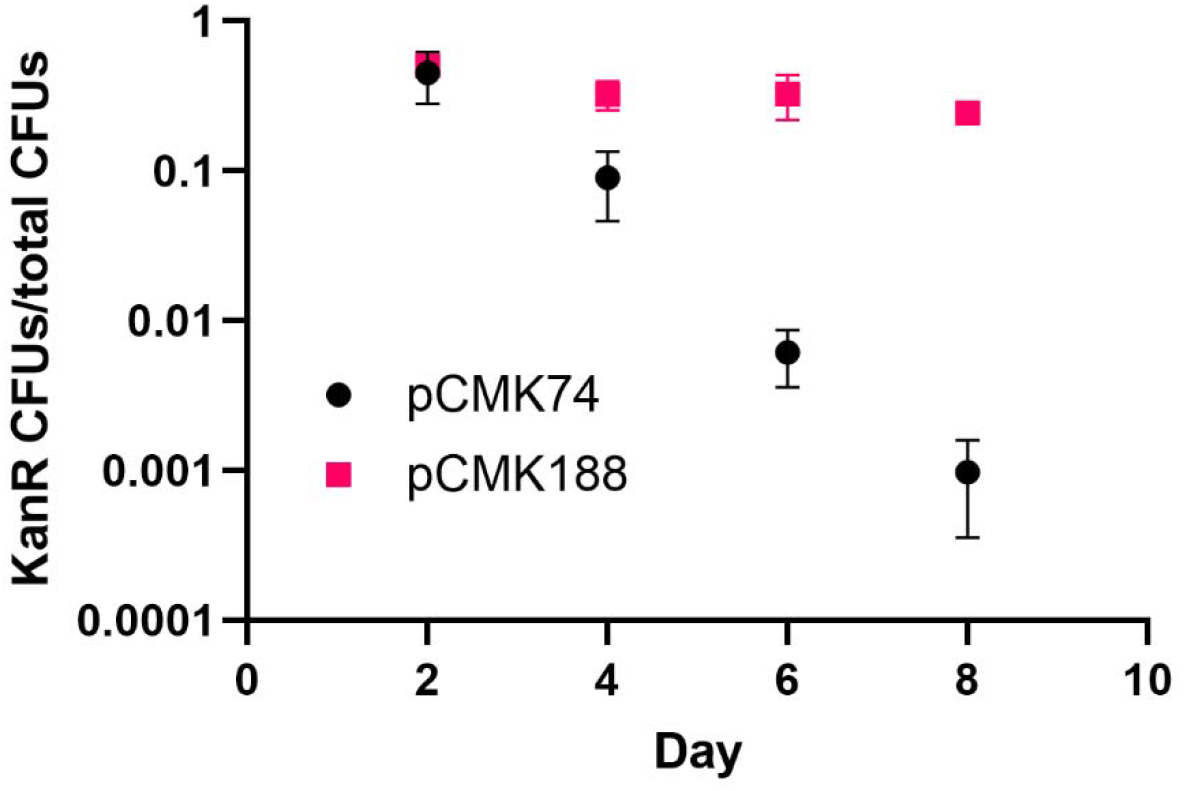
Stability of pCMK74 and pCMK188 in PCC 7002 when cultured in absence of antibiotic selection. Every two days, cultures were serially transferred into fresh media A+ without antibiotics. Error bars represent standard deviation from at least triplicate experiments.

### 3.4 Higher order plasmid multimers enhance natural transformation in PCC 7002

The above results were primarily obtained using plasmid samples miniprepped from *E. coli* and, as revealed by nanopore sequencing, naturally contained a small portion of plasmid multimers (data not shown). However, despite supporting successful transformations, direct use of miniprepped plasmid mixtures impedes our ability to decipher the impact of plasmid multimers on natural transformation efficiencies in PCC 7002. To facilitate a more detailed investigation of this relationship, an additional shuttle vector, pCMK139 (Figure 1), was constructed with the goal of enhancing total multimer synthesis in *E. coli*. In this case, the pBR322 origin in pCMK188 was replaced with a mutated pSC101 origin, where deletion of the par locus of pSC101 (responsible for resolving plasmid multimers back into monomers) has been shown to promote greater multimer accumulation [35]. Meanwhile, since pSC101 is a low copy number plasmid in *E. coli* (which limits the ability to prepare of large quantities of plasmid for fractionation by gel electrophoresis and extraction), the RepA gene of pSC101 (which governs plasmid copy number) was mutated (RepA^N99D^) to further increase both plasmid copy number and multimer yield [36]. Miniprepped samples of pCMK139 were separated via gel electrophoresis to resolve and extract individual fractions (Figure 3A) which, as characterized via nanopore sequencing, were each found to be uniquely enriched with respect differently sized multimers (Figure 3B). Specifically, Band 1 contained mostly monomeric pCMK139 (4838 bp) but also a small amount of dimer, Band 2 contained mostly dimeric pCMK139 (9676 bp), and Band 3 contained both dimeric and trimeric pCMK139 (14514 bp). Each isolated fraction was then individually transformed into PCC 7002, with the resulting transformation efficiencies compared in Figure 3C. In this case, Band 3 yielded an average of 3000 CFU per μg DNA transformed per mL cells, representing a 30-fold improvement.

**Figure 3.**
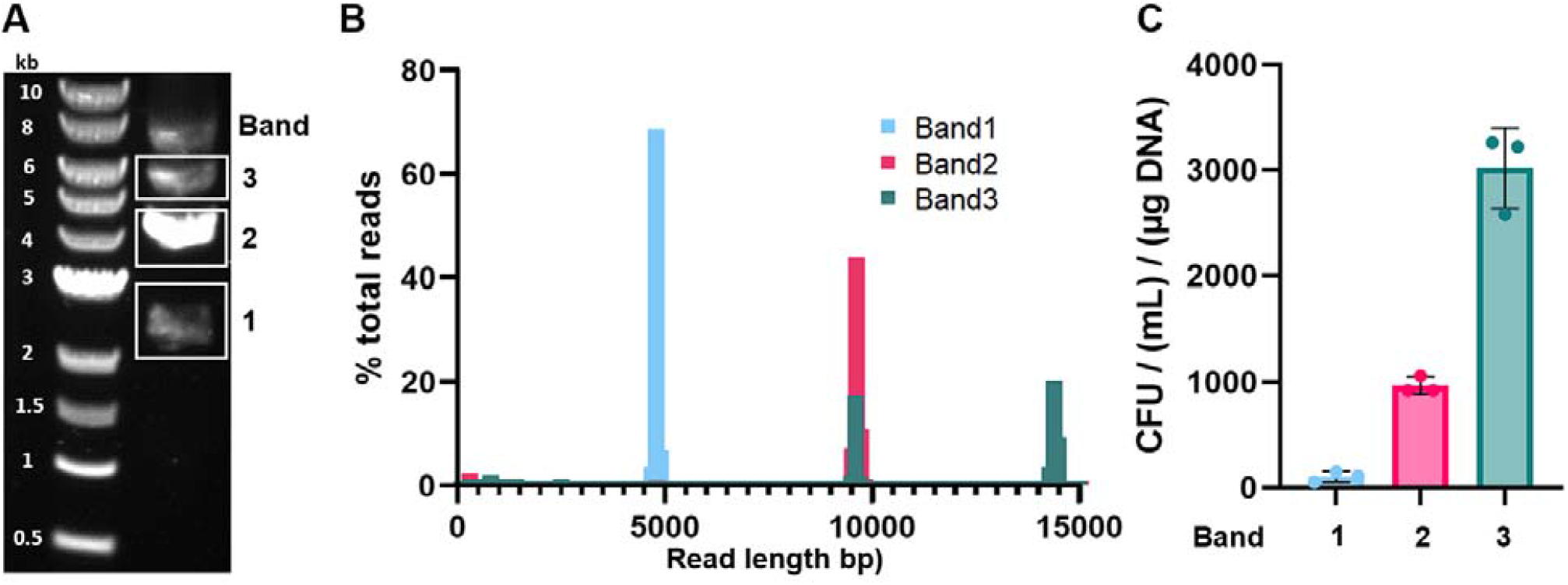
Isolation and transformation of different plasmid fractions uniquely enriched with different multimeric forms of pCMK139. A: Representative agarose gel showing how the original plasmid mixture was resolved via electrophoresis and the source of the three different plasmid fractions. The three fractions (i.e., major bands) were extracted, purified, and used to transform PCC 7002. B: Nanopore sequencing results obtained for each isolated plasmid fraction. Histograms compare the number of reads of a given length normalized by the total number of reads per sample, and were prepared using a bin size of 100 bp. Monomeric pCMK139 is 4838 bp in size, whereas its dimeric and trimeric forms are 9676 bp and 14514 bp, respectively. C: Effects of transforming different isolated plasmid fractions on transformation efficiency, as measured by the normalized colony forming units (CFUs) obtained for PCC 7002. CFUs were normalized by dividing by the volume of cells plated and mass of DNA transformed. Error bars represent standard deviation from triplicate experiments.

### 3.5 *E. coli* host selection influences multimer yields and transformation efficiency in PCC 7002

Since increased multimer content promotes enhanced transformation efficiencies, we lastly sought to determine if and how *E. coli* host selection may influence natural multimer abundance in miniprepped samples. While the above experiments solely utilized *E. coli* DHIP1 to prepare multimer containing samples via miniprep, its propensity for multimer synthesis was compared to that of DH5α and DH10β for both pCMK74 and pCMK188. As seen in Figure 4A, miniprepped samples from both DH10β and DHIP1 were enriched with multimers of pCMK188, but not so for DH5α. In contrast, pCMK74 samples contained only a minute fraction of dimeric plasmid regardless of *E. coli* strain used. Finally, it is noted that plasmids miniprepped from DHIP1 were also uniquely methylated at the included HIP1 site. Consistent with past studies in PCC 6803 [19], methylation was also found to improve transformation efficiencies for both plasmids in PCC 7002 (Figure 4B). For pCMK74, meanwhile, methylation had an outsized effect over multimer content, as no colonies were obtained for samples prepared from DH5α or DH10β. Meanwhile, plasmid multimers and plasmid methylation were both beneficial to but not essential for obtaining transformants of pCMK188.

**Figure 4.**
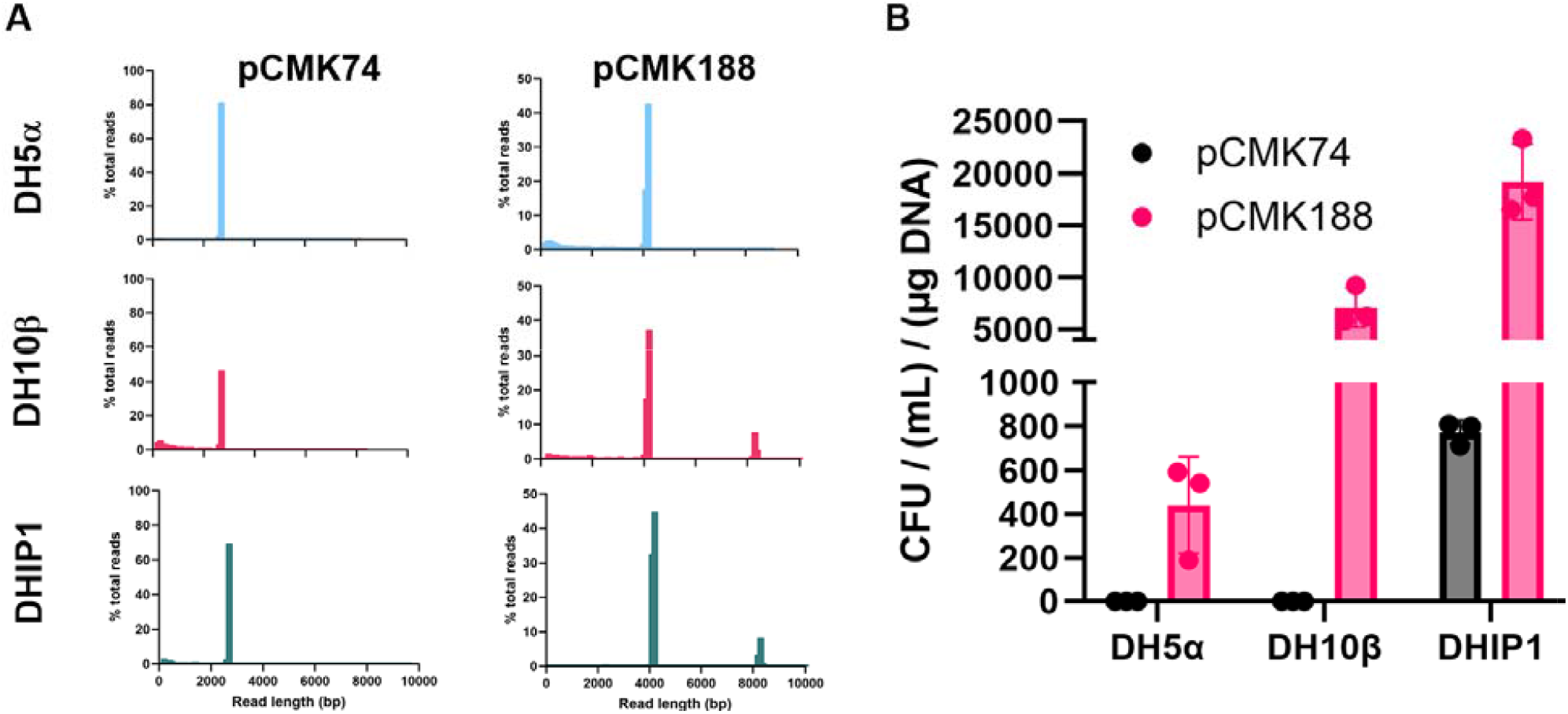
E. coli host selection influences multimer content and transformation efficiency of replicating plasmids in PCC 7002. A: Nanopore sequencing results obtained for pCMK74 and pCMK188 miniprepped from different E. coli strains. Histograms compare the number of reads of a given length normalized by the total number of reads per sample, and were prepared using a bin size of 100 bp. Monomeric pCMK74 is 2731 bp in size and its dimeric form is 5462 bp. Monomeric pCMK188 is 4195 bp in size and its dimeric form is 8390 bp. B: Transformation efficiencies of pCMK74 and pCMK188 when miniprepped from different E. coli hosts. CFUs were normalized by dividing by the volume of cells plated and amount of DNA transformed. Error bars represent standard deviation from triplicate experiments.

## 4. Discussion

To date, the most commonly employed replicating vectors in cyanobacteria include the RSF1010 broad host range plasmid [2,27,28] and shuttle vectors derived from native plasmids [4]. That said, RSF1010 is cumbersome to work with due to its large size (∼8.6 kb) and low copy number (just 10-12 copies per cell in *E. coli*[37]), along with the self-nicking function encoded by one of its mobilization proteins which further decreases plasmid yields [2]. To expand the portfolio of available and compatible plasmids for engineering PCC 7002, a set of novel replicating plasmids were herein developed based on the WVO1 broad host range origin and pCB2.4 from PCC 6803. To our knowledge, this represents the first report that the WVO1 origin stably replicates in any cyanobacterium. While WVO1-derived plasmids (pCMK74 and pCMK191) were maintained at a higher copy number than pCB2.4-derived plasmids (pCMK188 and pCMK139) in PCC 7002, the latter displayed much higher transformation efficiencies. Meanwhile, it is also interestingly noted that, in the absence of antibiotic selection, pCMK188 was more stable (i.e., resistant to curing) than pCMK74, perhaps due to the fact that pCB2.4 is derived from a cyanobacterium (PCC 6803) whereas WVO1 was originally discovered in *Streptococcus cremoris* [38]. Alternatively, given its low copy number, the greater persistence of pCMK188 could also be due to active segregation of plasmids between daughter cells [39]. Lastly, considering that it shares 96% genetic identity with PCC 7002, we furthermore expect that these novel plasmids will also be functional in the fast growing strain *Synechococcus* sp. PCC 11901 [40], and perhaps other strains as well.

Of the eight plasmids originally screened, four led to unhealthy colonies on plates that then quickly died upon transfer to liquid culture (Table 3). This could suggest that, although the associated origins might also be capable of replication in PCC 7002, they may not be stably maintained. Future engineering could perhaps address these unsuccessful outcomes, thereby providing additional cyanobacterial plasmid options. For example, instability may be a result of transcriptional read through into the origin of replication, which can cause a collision between RNA polymerase and plasmid replication machinery, decreasing plasmid copy number and stability [41,42].

Throughout this study, plasmids were miniprepped from *E. coli* and thereby naturally contained a small but important fraction of multimers. Since increased multimer content was found to correlate with enhanced transformation efficiencies (Figures 3 and 4), strategies for promoting multimer formation will facilitate future PCC 7002 engineering efforts. Multimer content in *E. coli*-derived plasmid samples is influenced by several factors, including host selection and the replication origin employed. For instance, although the underlying mechanism remains unclear, we found the multimer content to be greater for plasmids miniprepped from DH10β (and the derived strain DHIP1) than from DH5α. Both DH5α and DH10β contain the *recA1* mutation which, by suppressing plasmid recombination, has been shown to reduce multimer formation [13]. However, relative to DH5α, DH10β also contains numerous additional mutations, including gene duplications, multiple transposon insertions [43]. Future studies are needed to identify any specific genetic factors responsible for the observed difference. Meanwhile, with respect to the replication origin used, plasmids derived from WVO1 (pCMK74 and pCMK191) contained lower multimer content regardless of which *E. coli* strain they were miniprepped from. In both *E. coli* and cyanobacteria, the WVO1 origin notably replicates via rolling circle replication (RCR); a process that ultimately resolves generated multimeric plasmids back into their monomeric form [44]. Meanwhile, whereas pCB2.4-derived plasmids also replicate via RCR in cyanobacteria, only the pBR322 or pSC101 origins of pCMK188 and pCMK139 are active in *E. coli*; neither of which replicate via RCR but instead use theta replication [45]. Despite their reduced multimer content in *E. coli*, RCR plasmids may be ideally suited for natural transformation of cyanobacteria as they typically contain primosome assembly sites or a ssDNA origin of replication which, as described above, enables monomeric plasmids to be reformed upon entering the cell [46]. Taken together, we propose that future efforts to develop additional cyanobacterial replicating plasmids should prioritize engineering shuttle vectors that combine: 1) a cyanobacterial origin that replicates via RCR and 2) an *E. coli* origin that does not replicate by RCR or contains any other system for multimer resolution. Alternatively, multimeric plasmids can also be generated in vitro using prolonged overlap extension PCR (POE-PCR) [11]; an approach that might be particularly advantageous in cases where direct cloning into PCC 7002 (i.e., bypassing *E. coli*) is required.

## 5. Conclusion

A set of novel replicating plasmids were developed for *Synechococcus* sp. PCC 7002 based on the WVO1 and pCB2.4 origins of replication. The resulting plasmids were shown to be of stably maintained, including when co-transformed. Increased plasmid multimer content supports more efficient plasmid delivery via natural transformation and can be promoted via several easily implemented strategies.

## 6. Acknowledgements

This work was supported in part by a grant from the National Science Foundation (CBET-1705409). CMK received financial support in the form of a Completion Fellowship from Arizona State University. We thank Pranav Bhavaraju for his assistance with generating the Nanopore read length plots.

## Notes

### Competing Interest Statement

The authors have declared no competing interest.

